# Large-scale identification of viral quorum sensing systems reveals density-dependent sporulation-hijacking mechanisms in bacteriophages

**DOI:** 10.1101/2021.07.15.452460

**Authors:** Charles Bernard, Yanyan Li, Philippe Lopez, Eric Bapteste

## Abstract

Communication between viruses supported by quorum sensing systems (QSSs) were found to optimize the fitness of temperate bacteriophages of Bacilli by guiding the transition from the host-destructive lytic cycle to the host-protective lysogenic cycle in a density-dependent manner. All known phage-encoded QSSs consist of a communication propeptide and a cognate intracellular receptor that regulates the expression of adjacent target genes upon recognition of the matured peptide, a signature known as RRNPP and found in chromosomes, plasmids and phages of Firmicutes bacteria. Recently, we have introduced the RRNPP_detector software to detect novel genetic systems matching the RRNPP signature, which unearthed many novel phage-encoded candidate QSSs. Here, by looking at the adjacent genes likely regulated by these viral candidate QSSs, we identified an unsuspected clustering of viral QSSs with viral genes whose bacterial homologs are key regulators of the last-resort bacterial sporulation initiation pathway (*rap, spo0E* or *abrB*). Consistently, we found evidence in published data that certain of these QSSs encoded by prophages (phage genomes inserted within a bacterial genome) dynamically manipulate the timing of sporulation in the host. Because these viral QSSs are genetically diverse and are found associated with different sporulation regulators, this suggests a convergent evolution in bacteriophages of density-dependent sporulation-hijacking mechanisms.

**SIGNIFICANCE:** Communication between viruses supported by quorum sensing systems (QSSs) is a brand new research area that has transformed our views of viral adaptation and virus-host co-evolution. The viral QSSs discovered so far were found to guide the lysis-lysogeny decision in temperate bacteriophages as a function of phage density. Here, we identified that quorum sensing-mediated communication between phages can not only guide the regulation of viral processes but also the manipulation of the bacterial sporulation pathway. Our finding introduces the new view that not only bacteria decide when it is time to sporulate, some bacteriophages are also key stakeholders in this dynamical decision-making process. Considering that spores are the transmissive form of many pathogens, these new insights have important applied implications.

## INTRODUCTION

If cell-cell communication via quorum sensing was discovered in 1970 in bacteria, the first characterization of a functional viral quorum sensing systems (QSSs) only dates back to 2017 (1). Erez *et al*. discovered that certain Bacillus phages encode a communication propeptide, which upon expression, secretion, and maturation by the host cellular machinery, accumulates in the extracellular environment. Accordingly, during the lytic cycle, the extracellular concentration of this phage-encoded peptide reflects the number of host cells that have expressed the QSS-encoding genome of the phage, which correlates with the number of hosts that have been lysed by the phage (**Fig1**). At high concentrations of this peptide, when lots of hosts have been lysed and the survival of the phage-host collective can be endangered, the imported-peptide is transduced by its cognate phage-encoded intracellular receptor, which coordinates a population-wide transition from the host-destructive lytic cycle to the host-protective lysogenic cycle. Indeed, during the lysogenic cycle, the phage genome is inserted within the genome of the host, replicates as part of it, and often confers upon the host an immunity towards free virions This viral QSS, coined arbitrium, thereby optimizes the fitness of a phage with respect to its social context.

**Figure 1.**
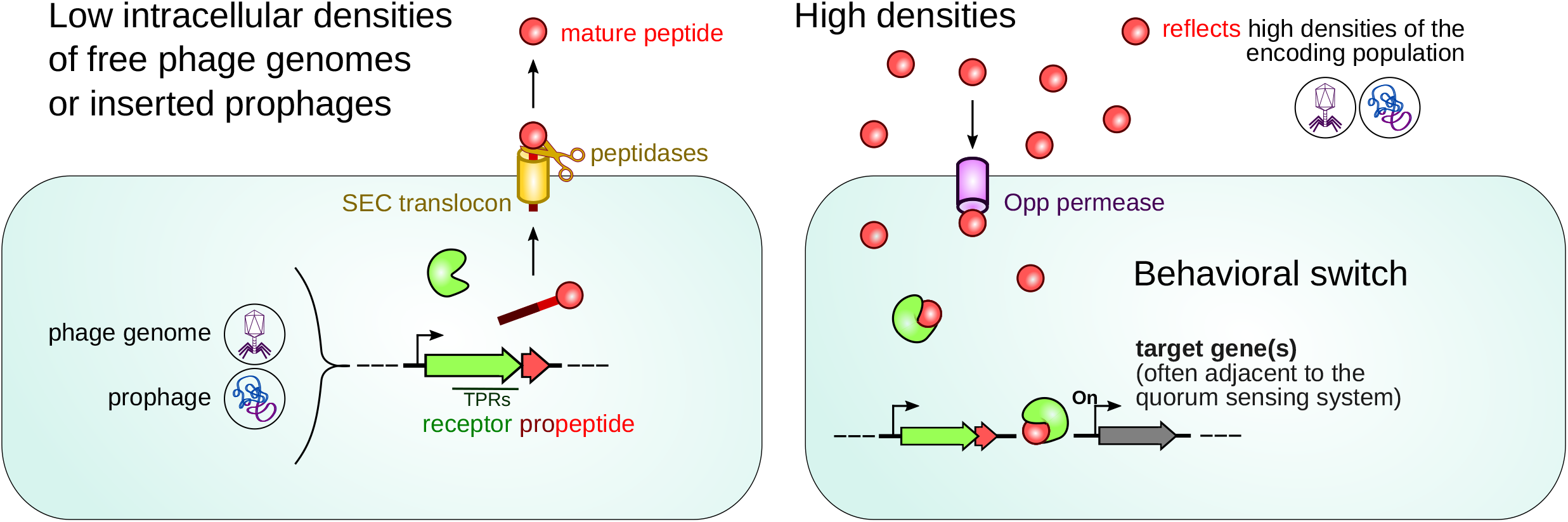
Mechanism of RRNPP QSSs in phages/prophages. The receptor and the communication propeptide of the phage-encoded QSS are in green and red, respectively. Upon bacterial expression, secretion and maturation, the concentration of the communication peptide reflects the intracellular density of phage genomes or prophages, which correlates with the number of lysed cells during the lytic cycle and the number of lysogenized cells during the lysogenic cycle. At high concentrations of the peptide, reflecting a quorum of phages/prophages, the receptor binds the imported mature peptide via TPRs motifs and gets either turned-on or -off as a transcription factor or a protein inhibitor, which is at the basis of density-dependent regulations of target genes (often adjacent to the QSS) or proteins. These regulations thereupon coordinates a behavioral transition at the scale of the entire phage/prophage population.

Arbitrium QSSs are classified as RRNPP QSSs, which consist of a communication propeptide (secreted and matured into a quorum sensing peptide) coupled with an intracellular receptor, turned-on or -off upon binding with the imported mature peptide (**Fig1**) (2). RRNPP receptors often are transcription factors that dynamically regulate the expression of adjacent genes. As genetically diverse as RRNPP QSSs are in chromosomes, plasmids and phages of Firmicutes, they however function according to the same canonical mechanism (**Fig1)**, except with very few exceptions in the mode of secretion (Shp and PrgQ peptides) (2). We recently identified that this shared mechanism between different RRNPP QSSs underlies a common signature that can be detected in-silico (3). On this basis, we have developed RRNPP_detector, a tool designed to identify novel candidate QSSs matching the RRNPP signature (3).

In a large-scale application of RRNPP_detector, we identified many novel candidate RRNPP QSSs in sequenced genomes of temperate bacteriophages and in prophages inserted within bacterial genomes (3). During the development of RRNPP_detector, we noticed an unsuspected clustering of viral candidate QSSs with adjacent viral genes whose bacterial homologs are key regulators of the bacterial sporulation initiation pathway. Because RRNPP QSSs tend to dynamically regulate adjacent genes (1, 2, 4, 5), this hinted at density-dependent manipulations of bacterial sporulation by phages.

In Firmicutes, sporulation leads to the formation of endospores, able to resist extreme environmental stresses for prolonged periods and to resume vegetative growth in response to favorable changes in environmental conditions (6). The sporulation pathway is initiated when transmembrane kinases sense stress stimuli, and thereupon transfer their phosphate, either directly (*Clostridium*) or via phosphorelay (*Bacillus, Brevibacillus*) to Spo0A, the master regulator of sporulation (7, 8). However, only high Spo0A-P concentrations, and therefore intense stresses, can commit a cell to sporulate (9). Because sporulation is costly, Spo0A-P accumulation is subjected to multiple regulative check points by the Rap, Spo0E and AbrB proteins (10, 11). In adverse circumstances, Rap, Spo0E and AbrB thereby form a decision-making regulatory circuit that controls the timing of sporulation in Bacilli (11).

Here, we present in more details our results regarding the association of sporulation regulators (either *rap, spo0E* or *abrB*) with candidate QSSs found in temperate bacteriophages of Firmicutes, their mechanistic consequence on the host biology, and their fundamental and applied implications.

## RESULTS AND DISCUSSION

### Identification of 384 viral candidate RRNPP QSSs, distributed into 26 families, of which only 6 were previously known

To detect viral QSSs, we followed the study design displayed in **FigS1**. We applied RRNPP_detector (15-65aa and 250-460aa length thresholds for propeptides and receptors) against the Gut Phage Database (12), 32,327 NCBI complete genomes of Viruses, and 3,577 NCBI complete genomes of Firmicutes (because phage genomes can be inserted within bacterial genomes). This led to the identification of 16 candidate RRNPP QSSs on intestinal phage genomes, 10 on sequenced genomes of temperate phages and 2671 on bacterial genomes, respectively. Prophage regions within bacterial genomes were subsequently detected by Phaster (13) and Prophage Hunter (14) to distinguish between genuine bacterial QSSs and prophage-encoded QSSs inserted within bacterial chromosomes or plasmids. This enabled identifying 358 additional candidate viral QSSs: 174 on intact/active prophages, 68 on questionable/ambiguous prophages, 116 on incomplete prophages. The genomic and taxonomic details of these 384 viral candidate RRNPP QSSs can be found in **TableS1**.

In a blast all vs all, the receptors of all the 2697 candidate QSSs were further classified into 64 groups of homologs / QSS families (sequence identity >=30%; mutual length coverage of the alignment >=80%), as in (3). Of these 64 QSS families, 26 families comprised at least one phage/prophage-encoded QSS and served as the focus of this study (metadata of the 26 families in **TableS1;** distribution of the 384 viral candidate RRNPP QSSs into these 26 families in **Fig2**). 6 of these 26 QSS families had already been described prior to our study: Rap-Npr of *Bacilli* (2, 15), AimR of *B. subtilis* phages (1), AloR of *Paenibacilli* (16), AimR of *B. cereus* phages (5), the family of QSSs characterized by Feng *et al*. in *Clostridium saccharoperbutylacetonicum* (17), the family of QSSs characterized by Kotte *et al*. in *Clostridium acetobutylicum* (18) (**Fig2**). Accordingly, the 20 other families show great promise to expand the known diversity of viral RRNPP QSSs. As expected, the arbitrium systems of *B. subtilis* (N=15) and *B. cereus* (N=9) phages form two 100% viral families (**Fig2B**). The two biggest families, Rap-Npr of Bacilli (N=2258) and the novel candidate QSS1 family of Brevibacilli (N=18) are found to be shared between bacterial chromosomes, plasmids and phages/prophages (**Fig2**). The Rap-Phr family notably is the family in which (pro)phage-encoded QSSs are the most prevalent (326 viral Rap-Phr QSSs (**TableS1 and Fig2A**)).

**Figure 2.**
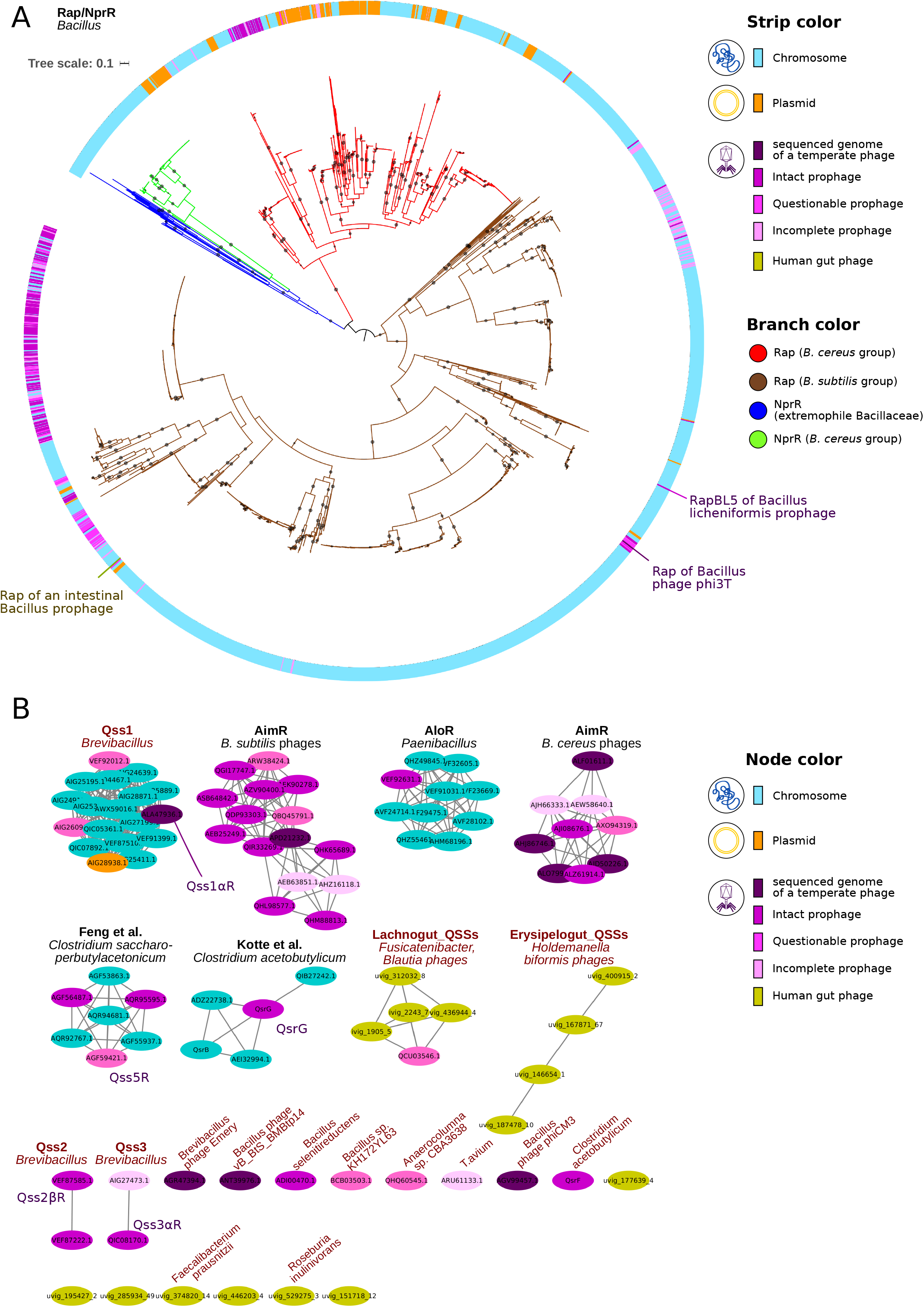
QSS families comprising at least one phage/prophage-encoded candidate QSS. **A. Maximum-likelihood phylogeny of receptors of the Rap/NprR family.** The figure displays the maximum-likelihood phylogenetic tree of the family comprising the Rap (no DNA binding domain) and the NprR (DNA binding domain) receptors that are part of a detected RRNPP-type QSS. The clustering of Rap and NprR into the same protein family is consistent with the common phylogenetic origin proposed for these receptors (S. Perchat, *et al*., *Microb. Cell* 2016)). The tree was midpoint rooted and a small black circle at the middle of a branch indicates that the branch is supported by 90% of the 1000 ultrafast bootstraps performed. Branch colors are indicative of the type of receptor (Rap or NprR) and of the bacterial group that either directly encodes the QSS or can be lysogenized by a (pro)phage that encodes the QSS. The colorstrip surrounding the phylogenetic tree assigns a color to each leaf based on the type of genetic support that encodes the QSS: blue for chromosomes, orange for plasmids, dark purple for sequenced genomes of temperate bacteriophages, different levels of purple for Phaster-predicted intact, questionable and incomplete prophages. The Rap receptors of Bacillus phage phi3T (only Rap found in a sequenced genome of a temperate phage) and of *B. licheniformis* intact prophage (viral Rap inserted into the host chromosome and shown to modulate the bacterial sporulation and competence pathways) are outlined. **B. Sequence similarity network of the other candidate QSS families comprising at least one phage/prophage-encoded QSS**. Each node corresponds to a receptor sequence found adjacent to a candidate pro-peptide and is colored according to the type of genetic element encoding the QSS, as displayed in the legend. The label of each node indicates the NCBI id or a Gut Phage Database id of the candidate receptor. Each edge corresponds to a similarity link between two receptors defined according to the following thresholds: percentage identity >= 30%, alignment coverage >= 80% of the lengths of both receptors, E-value <= 1E-5. Each connected component of the graph thereby defines groups of homologous receptors and is considered as a QSS family. The families are ordered from the largest to the smallest. Families with a black label were already described before (but not necessarily in phages) while families with a red label are novel. The most prevalent encoding-taxon in the family is displayed on top of it. The nodes of the receptors that are part of a predicted sporulation-hijacking QSS are characterized by an additional label.

### Rap-Phr QSSs that delay the timing of sporulation are found in many, diverse Bacillus bacteriophages

Importantly, bacterial Rap-Phr QSSs are known to regulate the competence, the sporulation and/or the production of public goods in *Bacilli*. Notably, these communication systems ensure that Spo0A-P only accumulates when the Rap-Phr encoding subpopulation reaches high densities (19). On this basis, Rap-Phr QSSs have been proposed as a means for a cell to delay a costly commitment to sporulation as long as the ratio of available food per kin-cell is compatible with individual survival in periods of nutrient limitation (19). However, the fact that Rap-Phr QSSs are found on phages implies that this delay in the timing of sporulation can sometimes be dependent on the intracellular density of phage genomes or prophages rather than on actual cell densities, presumably for the evolutionary benefit of the phage or the prophage-host collective (20).

Here, we inferred the maximum-likelihood phylogeny of the 2258 detected Rap receptors (**Fig2A**) and identified that the 326 viral Rap-Phr QSSs are genetically diverse and polyphyletic (scattered between bacterial leaves). Remarkably, this suggests that bacteria and phages frequently exchange these communication systems.

Indeed, from a phage’s perspective, a dynamical modulation of sporulation can be advantageous because on the one hand, sporulation can elicit the lytic cycle (21) and trigger cannibalistic behaviors that may reduce the amount of potential hosts (22) and on the other hand, spores can protect the phage genome under unfavorable environmental conditions. From a lysogenized host perspective, the *rap-phr* QSS acquired from the prophage can be considered as adaptive because they might support a cell-cell communication between lysogenized bacteria (since prophage density correlates with host density) that enables to cheat (delay production of public goods) and delay the costly sporulation program in a density-dependent manner, to the benefit of the prophage-host collective (23). A delay of sporulation might notably offer some additional time to resume growth if a peculiar stress happens to be relieved from the environment (24) or to benefit from the last bite of food present in the environment to replicate before sporulating, hence maximizing the number of representatives in the future heterogeneous population of spores.

### Multiple occurrences of the quorum sensing-mediated sporulation-hijacking genomic signature in temperate bacteriophages

Remarkably, the phage-encoded Rap-Phr QSSs were not the only potential host-hijacking viral QSSs. Indeed, we identified 5 additional candidate phage/prophage-encoded QSSs, distributed into 5 different QSSs families, predicted to manipulate the host sporulation initiation pathway in a density-dependent manner (**Fig3**). This prediction lies on the observation that their receptor harbors a DNA binding domain (**TableS1**) and thus likely regulates the expression of adjacent genes (like the arbitrium system), and that either the *spo0E* or *arbB* sporulation regulator is found in the vicinity of the QSSs on the phage/prophage DNA (**Fig3**). The same genomic context, albeit not encoded by a phage, was shown to be linked to sporulation regulation in *Paenibacillus* bacteria (16). By either activating or repressing the viral *spo0E* or *abrB* genes once a quorum of phages/prophages is reached, our candidate viral QSSs could influence the total concentration of Spo0E or AbrB within hosts in a density-dependent-manner, thus influencing the dynamics of Spo0A-P accumulation in these hosts and thereby modulating the sporulation initiation pathway to the benefit of the phage or the prophage (**FigS2**). Importantly, as the sporulation initiation pathway can trigger a wide range of biological processes (sporulation, biofilm formation, cannibalism, toxin production and solventogenesis (22, 25)), these viral candidate QSSs would also manipulate, in a density-dependent manner, a substantially broader spectrum of the host biology than spore formation alone (**FigS2**). Because these viral QSSs belong to distinct QSS families and are carried by phages infecting different hosts, this suggests a convergent evolution of density-dependent sporulation-hijacking in bacteriophages.

**Figure 3.**
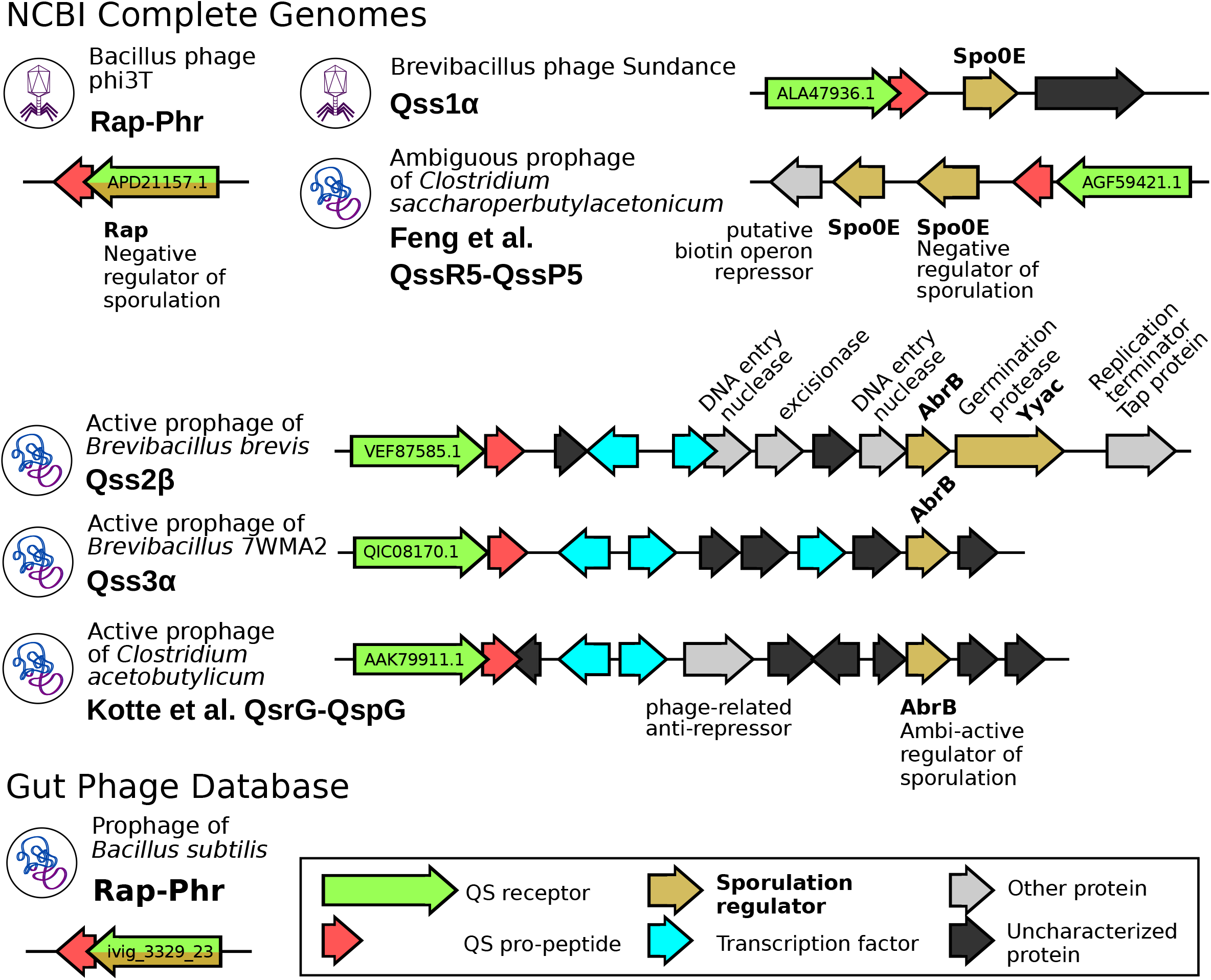
Clustering of sporulation initiation regulators with QSSs found in sequenced genomes of phages (virion icon) or prophages (lysogenized chromosome icon). The name and the genome of each QSS is written aside from each genomic context. Genes are colored according to their functional roles, as displayed in the legend. Within the QSS receptor gene, the NCBI id of the protein is shown. Because RRNPP QSSs tend to regulate adjacent genes, these genomic contexts hint at density-dependent sporulation-hijacking mechanisms.

We next found published data that support our prediction of a dynamical sporulation-hijacking mediated by phage-encoded QSSs. Indeed, the RapBL5-PhrBL5 of *Bacillus licheniformis* (26), the Qss5R-Qss5P of *Clostridium saccharoperbutylacetonicum (17)*, and the QsrG-QsrG of *Clostridium acetobutylicum* (18) were previously found to dynamically regulate sporulation and we identified that these QSSs are predicted by Prophage Hunter to be encoded by prophages inserted within bacterial chromosomes (**Fig4**). Moreover, 2 of these 3 prophages are predicted by Phaster to be intact and by Prophage Hunter to be active and thus able to re-initiate the lytic cycle upon excision while the other probably was domesticated by its host (27). These observations are the evidence that some genomes of temperate bacteriophages encode a communication system that guides the manipulation of bacterial sporulation.

**Figure 4.**
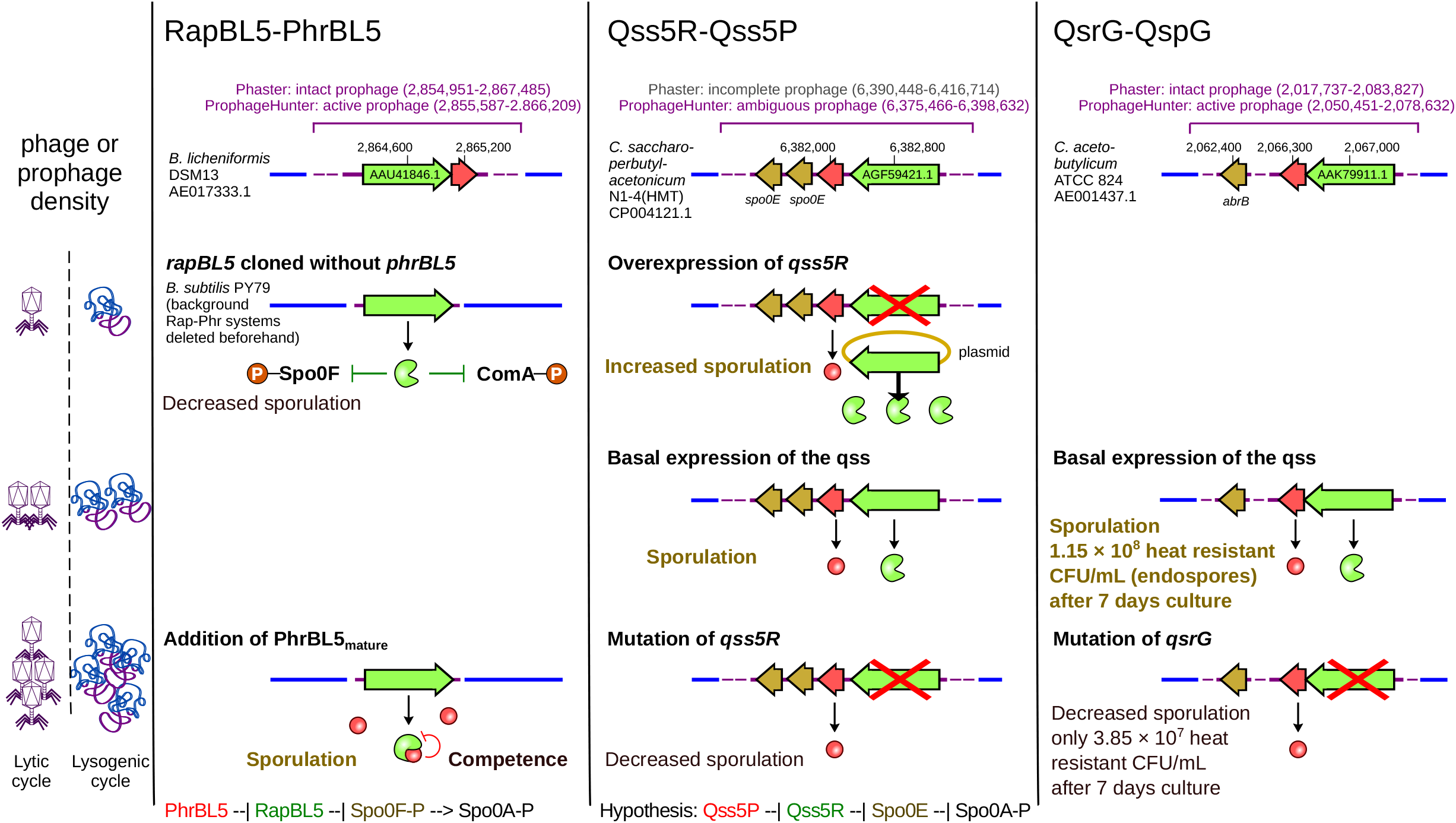
Summary of the evidences that some viral QSSs dynamically influence bacterial sporulation. The functions of the three QSSs (one column each) were investigated in (17, 18, 26) as displayed in this figure. The prediction that each QSS is encoded by a prophage is shown in purple at the top of each column. A prophage qualified as active by Prophage Hunter means that it is predicted to be able to reinitiate the lytic cycle upon excision from the bacterial chromosome. The interaction between the quorum sensing peptide was only investigated for RapBL5-PhrBL5 and is still lacking for Qss5R-Qss5P and QsrG-QspG. However, there is evidence in all QSSs that the receptor regulates the sporulation pathway, via Spo0F-P inhibition for the Rap protein inhibitor, and likely via transcriptional regulation of the adjacent *spo0E* and *abrB* genes for the Qss5R and QsrG transcription factors.

### Concluding remarks

Here, we computationally characterized distinct candidate QSSs presenting a genomic signature for a density-dependent hijacking of the bacterial sporulation initiation pathway in sequenced genomes of temperate bacteriophages (Bacillus phage phi3T, Brevibacillus phage Sundance) and in latent prophages. Moreover, we found published data that supports the validity of this signature. Because these QSSs are genetically diverse and belong to phages/prophages infecting different species of bacteria, this suggests multiple independent acquisitions of quorum sensing-mediated sporulation-hijacking genetic systems in phages. Accordingly, phage-encoded QSSs would not only dynamically regulate viral processes (as in the arbitrium QSSs) but also bacterial processes, for the benefit of the phage or the prophage-host collective. Our study also highlights that if bacteria decide when it’s time to sporulate, some phages also are stakeholders in the decision-making process. Some bacteriophages had already been reported to be either spore-restricting or spore-converting (28–31), but either way, these activations/impairments of sporulation were not the result of a decision making-process, unlike the density-dependent modulations described here.

From an applied viewpoint, these findings are also important because as sporulation enables bacteria to resist harsh environmental conditions, it represents a route for bacteria to travel between environments, and notably to end up within human bodies. Consequently, endospores are the transmissive form of many bacteria, be they commensal or pathogen (32–34). Furthermore, 50-60% of the bacterial genera from the human gut are estimated to produce resilient spores, specialized for host-to-host transmission (35). Accordingly, by dynamically interfering with sporulation, phage-encoded QSSs could influence the dynamics of transmission of bacteria in humans. With this respect, we also report a high prevalence of genes in the human gut phage database matching the HMM models of Spo0E and AbrB (**TableS2**), strengthening the case of a regulative effect of bacteriophages on the host-to-host transmission of gastrointestinal bacteria.

## METHODS

### Construction of the target data sets

The complete genomes of Viruses and Firmicutes were queried from the NCBI ‘Assembly’ database (36), as of 28/04/2020 and 10/04/2020, respectively. The features tables (annotations) and the encoded protein sequences of these genomes were downloaded using ‘GenBank’ as source database. The Gut Phage Database (12) was downloaded as of 29/10/2020, from the following url: http://ftp.ebi.ac.uk/pub/databases/metagenomics/genome_sets/gut_phage_database/

### Detection of candidate RRNPP QSSs

We launched a systematical search of the RRNPP-type signature using RRNPP_detector (3) against the complete genomes of Viruses and Firmicutes available on the NCBI and the MAGs of bacteriophages from the Gut Phage Database. RRNPP_detector defines candidate RRNPP-type quorum sensing systems as tandems of adjacent ORFs encoding a candidate receptor (protein matching HMMs of peptide-binding tetraticopeptide repeats (TPRs)) and a candidate pro-peptide (10-small protein predicted to be excreted via the SEC-translocon), consistent with the genetic features that are specific to the RRNPP quorum sensing mechanism and that are common between different experimentally-validated RRNPP QSSs (3). We specified a length range of 15-65aa for candidate propeptides and 250-460aa for candidate receptors.

### Classification of the candidate RRNPP QSSs into families

Because quorum sensing pro-peptides offer few amino acids to compare, are versatile and subjected to intragenic duplication (26), we classified the QSSs based on sequence homology of the receptors as in (3). We launched a BLASTp (37) All vs All of the receptors of the 2697 candidate QSSs identified in the complete genomes of Viruses and Firmicutes. The output of BLASTp was filtered to retain only the pairs of receptors giving rise to at least 30% sequence identity over more than 80% of the length of the two proteins. These pairs were used to build a sequence similarity network and the families were defined based on the connected components of the graph.

### Identification of already known QSS families

A BLASTp search was launched using as queries the RapA (NP_389125.1), NprR (WP_001187960.1), PlcR (WP_000542912.1), Rgg2 (WP_002990747.1), AimR (APD21232.1), AimR-like (AID50226.1), PrgX (WP_002366018.1), TraA (BAA11197.1), AloR13 (IMG: 2547357582), QsrB (AAK78305.1), Qss5R (AGF59421.1) reference receptors, and as a target database, the 2697 candidate QSS receptors previously identified. If the best hit of a reference RRNPP-type receptor gave rise to a sequence identity >= 30% over more than 80% mutual coverage, then the family to which this best hit belongs was considered as an already known family.

### Prophage detection

All the NCBI ids of the genomic accessions of chromosomes or plasmids of Firmicutes encoding one or several candidate QSSs were retrieved and automatically submitted to the Phaster webtool (13). Eventually, each QSS was defined as viral if its genomic coordinates on a given chromosome/plasmid fell within a region predicted by Phaster to belong to a prophage (qualified as either ‘intact’, ‘questionable’ or ‘incomplete’ prophage). Phaster results were complemented by ProphageHunter (14), a webtool that computes the likelihood that a prophage is active (able to reinitiate the lytic cycle by excision). Because ProphageHunter cannot be automatically queried, we only called upon this webtool for chromosomes/plasmids which encode QSSs that are not part of the two biggest of the 64 detected QSS families, namely Rap-Phr (2258 candidate QSSs) and PlcR-PapR (223 candidate QSSs, data not shown). Likewise, coordinates of candidate QSSs were eventually intersected with predicted prophage regions to detect potential prophage-encoded candidate QSSs that could have been missed by Phaster (**TableS1**). Based on the results of Phaster and Prophages, the focus of the study were further restricted to the 26 families comprising at least one phage or prophage-encoded QSSs.

### Prediction of the mature quorum sensing peptides

For each uncharacterized families of candidate receptors of size >1 with at least one (pro)phage-encoded member referenced in the NCBI, the cognate pro-peptides were aligned in a multiple sequence alignment (MSA) using MUSCLE *version 3*.*8*.*31* (38). Each MSA was visualized with Jalview *version 1*.*8*.*0_201* under the ClustalX color scheme which colors amino acids based on residue type conservation (39). The region of RRNPP-type pro-peptides encoding the mature quorum sensing peptide usually corresponds to a small sequence (5-6aa), located in the C-terminal of the pro-peptide, with conserved amino-acids types in at least 3 positions (1, 18, 20, 40). Based on the amino-acid profile of C-terminal residues in each MSA, putative mature quorum sensing peptides were manually determined (**FigS3**).

### Phylogenetic tree of Rap

A multiple sequence alignment (MSA) of the protein sequences of the Rap receptors forming a candidate Rap-Phr QSS was performed using MUSCLE *version 3*.*8*.*31* (38). The MSA was then trimmed using trimmal *version 1*.*4*.*rev2*2 with the option ‘-automated1’, optimized for maximum likelihood phylogenetic tree reconstruction (41). The trimmed MSA was then given as input to IQ-TREE *version multicore 1*.*6*.*10* to infer a maximum likelihood phylogenetic tree under the LG+G model with 1000 ultrafast bootstraps (42). The tree was further edited via the Interactive Tree Of Life (ITOL) online tool (43).

### Analysis of the genomic context of candidate QSSs

The genomic context of the (pro)phage-encoded candidate QSSs that are not part of the arbitrium families (functions already known in phages) and the Rap-Phr family (Rap is a protein inhibitor and not a transcription factor) and whose receptors matched the DNA binding domain profiles of RRNPP transcription factors (Hidden Markov Models PFAM PF01381, Superfamily SSF47413, SMART SM00530, CATH 1.10.260.40) were investigated by analyzing the functional annotation of their adjacent protein-coding genes, or when missing, by launching a “Conserved Domains” search within their sequence and a BLASTp search of their sequence against the NR (non-redundant) protein database maintained by the NCBI.

### Identification of *rap, spo0E* and *abrB* genes in the Gut Phage Database

With HMMER, we launched an HMM search of reference HMMs of Rap (PFAM PF18801), Spo0E (PFAM PF09388) and AbrB (SMART SM00966) against all the protein sequences predicted from the ORFs of the MAGs from the Gut Phage Database. The hits were retained only if they gave rise to an E-value < 1E-5.

## DECLARATIONS

### Availability of data and materials

All the NCBI or Gut Phage Database IDs of the proteins discussed in this manuscript are available in the supplementary tables.

### Authors’ Contribution

C.B, Y.L, E.B and P.L conceived the study. C.B performed the analyses. C.B, Y.L and E.B wrote the manuscript with input from all authors. All documents were edited and approved by all authors.

## Acknowledgments

We would like to thank Dr. A. K. Watson for critical reading and discussion.

## REFERENCES

1. Z. Erez, et al., Communication between viruses guides lysis-lysogeny decisions. Nature 541, 488–493 (2017).

2. M. B. Neiditch, G. C. Capodagli, G. Prehna, M. J. Federle, Genetic and Structural Analyses of RRNPP Intercellular Peptide Signaling of Gram-Positive Bacteria. Annu. Rev. Genet. 51, 311–333 (2017).

3. C. Bernard, Y. Li, E. Bapteste, P. Lopez, RRNPP_detector: a tool to detect RRNPP quorum sensing systems in chromosomes, plasmids and phages of gram-positive bacteria AUTHORS. bioRxiv, 2021.08.18.456871 (2021).

4. V. Kohler, W. Keller, E. Grohmann, Regulation of gram-positive conjugation. Front. Microbiol. 10, 1134 (2019).

5. A. Stokar-Avihail, N. Tal, Z. Erez, A. Lopatina, R. Sorek, Widespread Utilization of Peptide Communication in Phages Infecting Soil and Pathogenic Bacteria. Cell Host Microbe 25, 746–755.e5 (2019).

6. M. Y. Galperin, Genome Diversity of Spore-Forming Firmicutes. Microbiol. Spectr. 1, TBS–0015-2012 (2013).

7. I. S. Tan, K. S. Ramamurthi, Spore formation in Bacillus subtilis. Environ. Microbiol. Rep. 6, 212–225 (2014).

8. M. A. Al-Hinai, S. W. Jones, E. T. Papoutsakis, The Clostridium Sporulation Programs: Diversity and Preservation of Endospore Differentiation. Microbiol. Mol. Biol. Rev. 79, 19–37 (2015).

9. M. Fujita, R. Losick, Evidence that entry into sporulation in Bacillus subtilis is governed by a gradual increase in the level and activity of the master regulator Spo0A. Genes Dev. 19, 2236–2244 (2005).

10. S. H. Shafikhani, T. Leighton, AbrB and Spo0E Control the Proper Timing of Sporulation in Bacillus subtilis. Curr. Microbiol. 48, 262–269 (2004).

11. D. Schultz, M. Lu, T. Stavropoulos, J. Onuchic, E. Ben-Jacob, Turning oscillations into opportunities: Lessons from a bacterial decision gate. Sci. Rep. 3 (2013).

12. L. F. Camarillo-Guerrero, A. Almeida, G. Rangel-Pineros, R. D. Finn, T. D. Lawley, Massive expansion of human gut bacteriophage diversity. Cell 184, 1098–1109.e9 (2021).

13. D. Arndt, et al., PHASTER: a better, faster version of the PHAST phage search tool. Nucleic Acids Res. 44, W16 (2016).

14. W. Song, et al., Prophage Hunter: an integrative hunting tool for active prophages. Nucleic Acids Res. 47, W74–W80 (2019).

15. S. Perchat, et al., NprR, a moonlighting quorum sensor shifting from a phosphatase activity to a transcriptional activator. Microb. Cell 3, 573–575 (2016).

16. M. Voichek, S. Maaß, T. Kroniger, D. Becher, R. Sorek, Peptide-based quorum sensing systems in Paenibacillus polymyxa. Life Sci. Alliance 3 (2020).

17. J. Feng, et al., RRNPP-Type quorum-sensing systems regulate solvent formation, sporulation and cell motility in Clostridium saccharoperbutylacetonicum. Biotechnol. Biofuels 13, 1–16 (2020).

18. A. K. Kotte, et al., RRNPP-type quorum sensing affects solvent formation and sporulation in clostridium acetobutylicum. Microbiol. (United Kingdom) 166, 579–592 (2020).

19. I. B. Bischofs, J. A. Hug, A. W. Liu, D. M. Wolf, A. P. Arkin, Complexity in bacterial cell-cell communication: quorum signal integration and subpopulation signaling in the Bacillus subtilis phosphorelay. Proc. Natl. Acad. Sci. U. S. A. 106, 6459–64 (2009).

20. C. Bernard, Y. Li, P. Lopez, E. Bapteste, Beyond arbitrium: identification of a second communication system in Bacillus phage phi3T that may regulate host defense mechanisms. ISME J. (2020) https://doi.org/10.1038/s41396-020-00795-9.

21. W. J. Meijer, et al., Molecular basis for the exploitation of spore formation as survival mechanism by virulent phage φ29. EMBO J. 24, 3647–3657 (2005).

22. J. E. González-Pastor, Cannibalism: A social behavior in sporulating Bacillus subtilis. FEMS Microbiol. Rev. 35, 415–424 (2011).

23. M. Kalamara, M. Spacapan, I. Mandic-Mulec, N. R. Stanley-Wall, Social behaviours by Bacillus subtilis: quorum sensing, kin discrimination and beyond. Mol. Microbiol. 110, 863 (2018).

24. J. E. González-Pastor, E. C. Hobbs, R. Losick, Cannibalism by Sporulating Bacteria. Science (80-.). 301, 510–513 (2003).

25. P. Dürre, et al., Transcriptional regulation of solventogenesis in Clostridium acetobutylicum in Journal of Molecular Microbiology and Biotechnology, (2002), pp. 295–300.

26. E. Even-Tov, S. Omer Bendori, S. Pollak, A. Eldar, Transient Duplication-Dependent Divergence and Horizontal Transfer Underlie the Evolutionary Dynamics of Bacterial Cell–Cell Signaling. PLOS Biol. 14, e2000330 (2016).

27. L. M. Bobay, M. Touchon, E. P. C. Rocha, Pervasive domestication of defective prophages by bacteria. Proc. Natl. Acad. Sci. U. S. A. 111, 12127–12132 (2014).

28. D. P. Boudreaux, V. R. Srinivasan, “Bacteriophage-induced Sporulation in Bacillus cereus T.”

29. M. G. Bramucci, K. M. Keggins, P. S. Lovett, Bacteriophage conversion of spore-negative mutants to spore-positive in Bacillus pumilus. J. Virol. 22, 194–202 (1977).

30. T. H. Silver-Mysliwiec, M. G. Bramucci, Bacteriophage-enhanced sporulation: Comparison of spore-converting bacteriophages PMB12 and SP10. J. Bacteriol. 172, 1948–1953 (1990).

31. R. Schuch, V. A. Fischetti, The secret life of the anthrax agent Bacillus anthracis: bacteriophage-mediated ecological adaptations. PLoS One 4, e6532 (2009).

32. M. Mallozzi, V. K. Viswanathan, G. Vedantam, Spore-forming Bacilli and Clostridia in human disease. Future Microbiol. 5, 1109–1123 (2010).

33. F. Postollec, et al., Tracking spore-forming bacteria in food: From natural biodiversity to selection by processes. Int. J. Food Microbiol. 158, 1–8 (2012).

34. M. C. Swick, T. M. Koehler, A. Driks, Surviving Between Hosts: Sporulation and Transmission. Microbiol. Spectr. 4 (2016).

35. H. P. Browne, et al., Culturing of “unculturable” human microbiota reveals novel taxa and extensive sporulation. Nature 533, 543–546 (2016).

36. NCBI Resource Coordinators, Database resources of the National Center for Biotechnology Information. Nucleic Acids Res. 44, D7–D19 (2016).

37. S. F. Altschul, W. Gish, W. Miller, E. W. Myers, D. J. Lipman, Basic local alignment search tool. J. Mol. Biol. 215, 403–410 (1990).

38. R. C. Edgar, MUSCLE: multiple sequence alignment with high accuracy and high throughput. Nucleic Acids Res. 32, 1792–7 (2004).

39. A. M. Waterhouse, J. B. Procter, D. M. A. Martin, M. Clamp, G. J. Barton, Jalview Version 2--a multiple sequence alignment editor and analysis workbench. Bioinformatics 25, 1189–1191 (2009).

40. M. Pottathil, B. A. Lazazzera, The extracellular PHR peptide-Rap phosphatase signaling circuit of bacillus subtilis. Front. Biosci. 8, 913 (2003).

41. S. Capella-Gutierrez, J. M. Silla-Martinez, T. Gabaldon, trimAl: a tool for automated alignment trimming in large-scale phylogenetic analyses. Bioinformatics 25, 1972–1973 (2009).

42. L.-T. Nguyen, H. A. Schmidt, A. von Haeseler, B. Q. Minh, IQ-TREE: A Fast and Effective Stochastic Algorithm for Estimating Maximum-Likelihood Phylogenies. Mol. Biol. Evol. 32, 268–274 (2015).

43. I. Letunic, P. Bork, Interactive Tree of Life (iTOL) v4: Recent updates and new developments. Nucleic Acids Res. 47, W256–W259 (2019).

