## Supplementary figures for "Large-scale identification of viral quorum sensing systems reveals density-dependent sporulation-hijacking mechanisms in bacteriophages"

**Figure S1: Study design**.

RRNPP-detector was applied against complete genomes of Firmicutes and Viruses from the NCBI and against the Gut Phage Database. This algorithm defines candidate RRNPP QSSs as tandems of adjacent ORFs encoding a candidate receptor (250-460aa protein matching HMMs of peptide-binding tetraticopeptide repeats (TPRs)) and a candidate pro-peptide (15-65aa protein predicted to be excreted via the SEC-translocon by SignalP). The candidate QSSs were further classified into families, defined as groups of homologous receptors in a BLASTp all vs all. This study further focused on families in which at least one QSS is encoded by a phage or a genomic region predicted by Phaster and/or ProphageHunter to belong to a prophage inserted within a bacterial genome. Subsequently, each QSS family with viral representatives was computationally characterized. Protein families of receptors shared between bacterial genomes and phage genomes were aligned, trimmed and given as input to IQ-TREE to construct phylogenetic trees in order to visualize if and how QSSs travel onto different kinds of genetic supports (chromosomes, plasmids, phage genomes) rather than to stay in their hosts lineages. In QSS families comprising more than one QSS, the propeptides were also aligned and visualized with Jalview to predict the sequence of each mature peptide. Finally, as RRNPP-type receptors that are transcription factors tend to regulate adjacent genes, the genomic neighborhood of each (pro)phage-encoded receptor with a detected DNA binding domain was analyzed to predict the functions regulated by the QSS.

**Figure S2: Predicted modulations of the sporulation initiation pathway mediated by phage/prophage-encoded QSSs**

The left and the right panels display the sporulation initiation pathway of the *Bacillus* genus and of the *Clostridium* genus, respectively. Transcriptional regulations are depicted by plain lines, whereas regulations at protein levels are depicted by dashed lines. At the end of each line, an arrow depicts an activation, a “T” symbol depicts an inhibition while a circle depicts an unknown direction of regulation. Written in grey are the inactive forms of sporulation proteins whereas their active, phosphorylated forms are written in black. The gradient of concentration starting from Spo0A-P indicates that sporulation is triggered by high levels of the master Spo0A-P regulator. Lower concentrations of Spo0A-P can trigger other bacterial processes than sporulation, as they may relieve a specific environmental stress and thus prevent, through alleviation of Spo0A phosphorylation, from a costly commitment to spore formation. The brown proteins (Rap, Spo0E and AbrB) depict regulators of Spo0A-P accumulation that are encoded by both bacteria and (pro)phages. The expression of (pro)phage-encoded *rap*, *spo0E* or *abrB* thus likely amplifies, by additive effect, the step of the host pathway controlled by each corresponding bacterial homolog. Red and green proteins depict the mature peptide and the receptor of a (pro)phage-encoded QSS inferred to regulate (pro)phage-encoded Rap, Spo0E or AbrB. An icon of grouped phages signifies that the regulation from the mature peptide to the receptor is expected to happen only at high (pro)phage densities. Each icon has its own color to highlight that the QSS genetic systems are encoded by different (pro)phages. These mechanisms are proposed to enable some bacteriophages to modulate the host sporulation initiation pathway and the competence pathway in a density-dependent manner.

**Figure S3: Multiple sequence alignments of QSS families cognate pro-peptides**

The figure displays the multiple sequence alignment of cognate propeptides for each receptor family of size > 1 that includes at least one (pro)phage-encoded QSS. A purple circle at the left of each protein identifier of the propeptide indicates that the QSS was found in a phage genome whereas a purple circle indicates that the QSS was found in a predicted prophage region. The residues are colored according to the ClustalX colorscheme (**http://www.jalview.org/help/html/colourSchemes/clustal.html**), which colors amino acids based on residue type conservation (hydrophobic, positively charged, negatively charged, polar etc…). Pro-peptides are characterized by a N-terminal region composed of positively charged amino acids (R, K), followed by a hydrophobic region. The mature peptide (typically 5 to 6 aminoacids) is usually encoded by a C-terminal region of the propeptide and is characterized in the alignment by the entanglement of conserved and variable positions.
